# Viscous shear is a key force in *Drosophila* ventral furrow morphogenesis

**DOI:** 10.1101/2021.04.21.440835

**Authors:** Amanda Nicole Goldner, Mohamad Ibrahim Cheikh, Miriam Osterfield, Konstantin Doubrovinski

## Abstract

Ventral furrow (VF) formation in *Drosophila melanogaster* is an important model of epithelial folding. Previous models of VF formation require cell volume conservation to convert apically localized constriction forces into lateral cell elongation and tissue folding. Here, we investigated embryonic morphogenesis in anillin knockdown (*scra* RNAi) embryos, where basal cell membranes fail to form and therefore cells can lose cytoplasmic volume through their basal side. Surprisingly, the mesoderm elongation and subsequent folding that comprise VF formation occurred essentially normally. We hypothesized that the effects of viscous shear may be sufficient to drive membrane elongation, providing effective volume conservation, and thus driving tissue folding. Since this hypothesis may not be possible to test experimentally, we turned to a computational approach. A minimal model of VF formation accounting for fluid dynamics indicated that shear forces can indeed explain our experimental observation. However, this conclusion depended on specific values of the model parameters. To test whether viscous shear is a dominant force for morphogenesis *in vivo*, we developed a highly realistic computational model incorporating both accurate cell and tissue geometry and experimentally measured material parameters. Results from this model demonstrate that viscous shear generates sufficient force to drive cell elongation and tissue folding *in vivo*.

## INTRODUCTION

Epithelial folding occurs throughout many stages in animal development (Leptin, 1991, Martinez Arias and Stewart, 2003), including neurulation (Maruyama and Andrew, 2012) and lung bud formation (Kim et al., 2013). During *Drosophila* gastrulation, the initially-monolayered embryo becomes multilayered through epithelial folding (Campos-Ortega and Hartenstein, 1985, Kohler, 1994).

Epithelial folding is essentially a physical process and therefore, comprehensive mechanistic understanding must include a knowledge of the mechanical processes involved. A common physical mechanism underlying many instances of epithelial folding is apical constriction (Martin and Goldstein, 2014, Sawyer et al., 2010). However, lateral and basal portions of the cell have previously been shown to influence epithelial folding (Gracia et al., 2019, Gutzman et al., 2018, Sui et al., 2018, Sui and Dahmann, 2020), although these effects are much less studied.

Likewise, the role of active tensions generated by the cytoskeleton have been examined in many systems (Heer and Martin, 2017, Quintin et al., 2008). However, although mechanical effects such as membrane elasticity, cytoplasmic viscosity and fluid flow have been implicated in some epithelial morphogenetic events (He et al., 2014, Luu et al., 2011, Mongera et al., 2018, Smutny et al., 2017), these effects have also been understudied.

*Drosophila* gastrulation is an ideal model system to address the physical mechanisms underlying epithelial folding, since it has been highly characterized genetically and morphologically (Barrett et al., 1997, Costa et al., 1994, Kanesaki et al., 2013, Kerridge et al., 2016, Kolsch et al., 2007, Lye and Sanson, 2011, Manning et al., 2013, Sweeton et al., 1991, Turner and Mahowald, 1977). In this work we examine the roles of basal membranes and cytoplasmic shear in ventral furrow (VF) formation during gastrulation. We first begin by reviewing the developmental processes that precede and set the stage for gastrulation in the fruit fly.

After fertilization of a *Drosophila* egg, the nuclei undergo 13 division cycles and migrate to the periphery of the embryo. Since nuclear division at this stage is not accompanied by cytokinesis, this results in an embryo that is essentially a large multinucleate cell encased in a rigid vitelline membrane (Leptin, 1991, Zalokar and Erk, 1976). Subsequently, lateral membranes grow inwards to compartmentalize peripheral nuclei into individual cells, forming the epithelium in a process known as cellularization (Campos-Ortega and Hartenstein, 1985, Lecuit and Wieschaus, 2000). Throughout the course of cellularization, cells remain open to the yolk sac such that there is a “hole” on the basal side of each cell (see schematic in Figure S1) (Krueger et al., 2019, Loncar and Singer, 1995, Mazumdar and Mazumdar, 2002).

As cellularization completes, gastrulation begins with the folding of the ventral furrow (VF) (Campos-Ortega and Hartenstein, 1985, Leptin and Grunewald, 1990, Sweeton et al., 1991, Turner and Mahowald, 1977). VF formation involves a sequence of shape changes in a subset of ventral cells, starting with constriction of their apical membranes (Kam et al., 1991, Leptin, 1991). As VF formation proceeds, lateral membranes lengthen before eventually shortening again as the basal membranes seal and the ventral tissue invaginates to form a furrow (Lye and Sanson, 2011, Sweeton et al., 1991, Turner and Mahowald, 1977). Initially, these shape changes were thought to be driven primarily by apical acto-myosin constriction (Martin et al., 2010, Martin et al., 2009, Sawyer et al., 2010). More recent work has indicated that lateral tensions may also play an important role (Gracia et al., 2019). Since cellularization is still ongoing during the initial apical constriction phase of VF formation, cells remain open and only start sealing basally as tissue invagination begins. The simultaneous occurrences of basal membrane formation and epithelial folding suggest that one process may rely on the other.

Here we began by exploring the functional importance of basal membrane formation to VF folding. In this study we show that basal membranes are not required for epithelial folding in the VF. We computationally explored alternative mechanisms for folding and found that viscous shear can compensate for the lack of basal membranes to maintain cell volume conservation and drive tissue folding. We then used a biophysically realistic model with experimentally measured parameters to test this model quantitatively, and found that this mechanism indeed operates in the time-scales and material property parameter regimes found *in vivo*.

## RESULTS

### Basal membranes are not required for VF formation

We began by exploring the role of basal membranes in VF formation. Loss-of-function mutations in Anillin (Scraps) – a component of the contractile ring scaffold that narrows to seal off cells and form basal membranes – delay basal membrane formation (Field et al., 2005, Thomas and Wieschaus, 2004, Xue and Sokac, 2016). However, gastrulation has not been characterized in this genetic background. For experimental simplicity, we first replicated this phenotype via maternal GAL4-driven expression of short hairpin RNA against *scra* (UAS- scraTRiP). Live imaging of embryos expressing maternal GAL4, UAS-scraTRiP, and a membrane marker (UAS-Nrt-GFP) showed that the effects of *scra* RNAi during cellularization were indistinguishable from those seen in Anillin loss-of-function mutants (Figure S2) (Field et al., 2005, Thomas and Wieschaus, 2004, Xue and Sokac, 2016). To better characterize gastrulation in this background, we prepared sections of heat-fixed gastrulas and stained them for Neurotactin (Nrt; a membrane marker), Snail (Sna; a mesoderm marker), and DAPI to assess morphology (Figure 1). As a control, embryos of similar genetic background without UAS- scraTRiP were stained according to the same protocol (Figures 1A-C). Although basal membranes never form in *scra* RNAi gastrulas, the VF consistently folds to its usual depth (Figures 1G-I).

**Figure 1.**
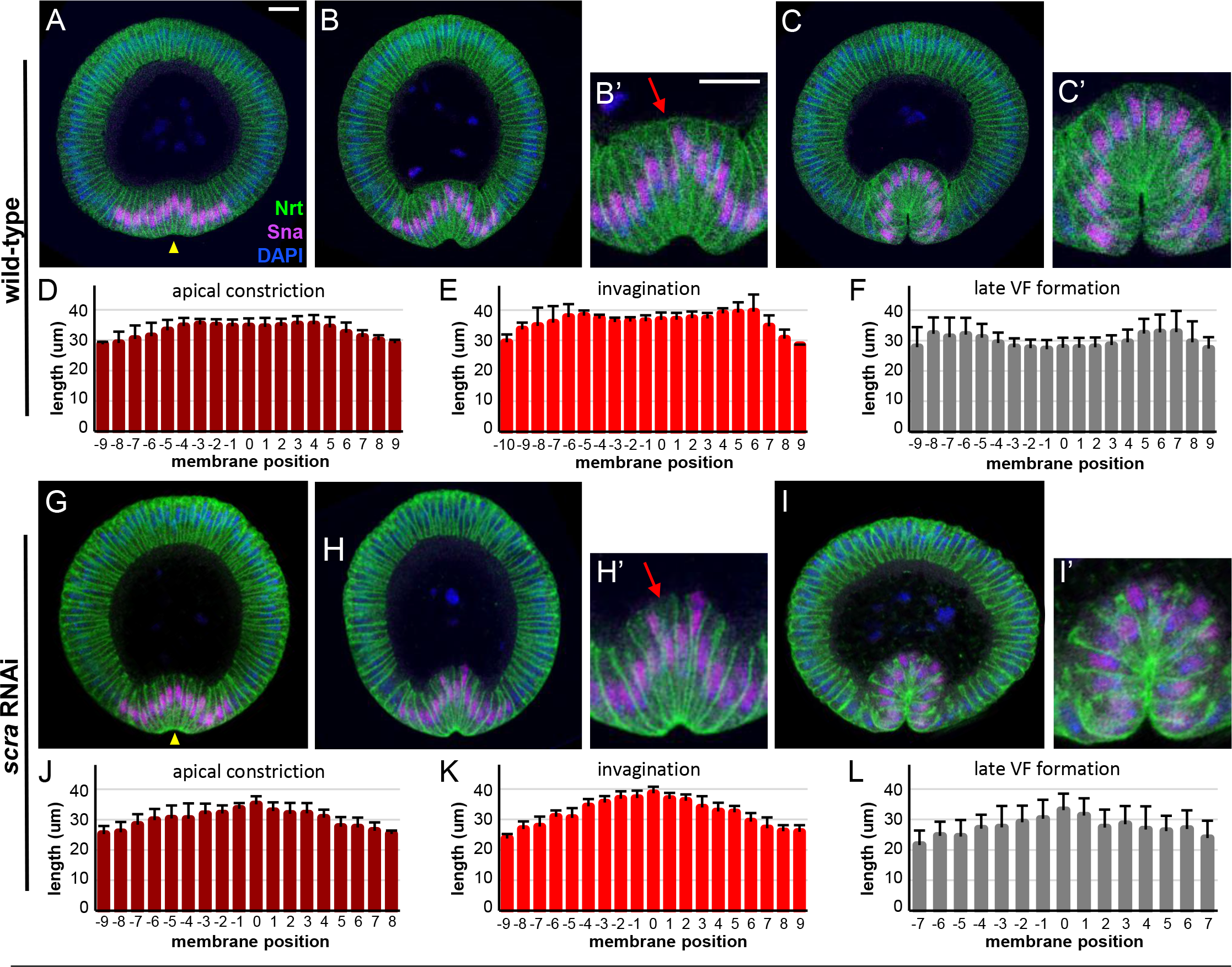
Basal membranes are not required for VF formation. Confocal immunofluorescence (A-C, G-I) and quantification of tissue morphology (D-F,J-L) in wild-type (A-F) and *scra* RNAi (G-L) embryos. (A,G) Embryos at apical constriction stage. Yellow arrowheads indicate apical surface of ventral mesoderm cells, which are constricting in both genetic backgrounds. (B,H) Embryos at early invagination stage. Ventrally-located (Snail-positive) mesoderm cells elongate along their apical-basal axis in both genotypes, and tissue invagination has begun. Basal surfaces, indicated by red arrows in (B’) and (H’), are closed in wild-type but remain open in *scra* RNAi embryos, (C,I) Embryos at late VF stage. Mesodermal cells have fully invaginated into the interior of the embryo in both genotypes, although basal surfaces still remain open in *scra* RNAi embryos. All images shown were cropped, rotated ventral side-down, and set against a black background. All scale bars are 20 μm. (D-F) Mesoderm lateral membrane lengths in wild-type embryos during (D) apical constriction (n = 9 embryos), (E) invagination (n = 7 embryos), and (F) late VF formation (n = 10 embryos). (J-L) Mesoderm lateral membrane lengths in *scra* RNAi embryos during (J) apical constriction (n = 8 embryos), (K) invagination (n = 8 embryos), and (L) late VF formation (n = 6 embryos). (B’), (C’), (H’), and (I’) are 2x magnifications of (B), (C), (H), and (I), respectively. Data are represented as mean ± s.d. Although wild-type and *scra* RNAi embryos appear qualitatively similar at the level of gross tissue morphology, lateral membrane lengths at each of the three stages of VF formation are significantly different between these two genotypes (wt vs. *scra* RNAi, p<0.001). See also Figures S2-S5.

### Membrane lengths are altered in *scra* RNAi embryos

We were interested to see if there were any morphological differences between *scra* RNAi and control embryos. To this end, we compared the distribution of lateral membrane lengths along transverse sections of control (Figures 1D-F) and *scra* RNAi (Figures 1J-L) embryos at various stages of VF formation, first confining our comparisons to mesoderm (Snail-positive) cells. In both genetic backgrounds, lateral membranes lengthen as apical constriction proceeds into invagination, then shorten as the VF forms more fully (Figures 1D-F, 1J-L). However, the spatial pattern of membrane lengths diverges in these genetic backgrounds as the VF forms. During early invagination, the peripheral mesoderm cells lengthen, while the equivalent *scra* RNAi cells remain essentially unchanged in length (Figure S3). Later, as the VF fully forms, the longest lateral membranes are located in the center of the furrow in *scra* RNAi embryos (single peak in Figure 1L), while the longest membranes in the control embryos are located peripherally within the VF (two peaks in Figure 1F). In particular, the membranes in the peripheral portions of the VF are strikingly shorter in *scra* RNAi embryos than in controls (Figures 1C,F,I,L and Figure S6A).

We also measured the length of ectodermal membranes (outside the VF) at the same three stages of VF formation (Figure S4). In both cases, ectodermal membranes lengthen during VF formation. However, ectodermal lengthening is delayed in *scra* RNAi embryos (Figure S4).

### *scra* RNAi VFs remain folded despite degradation of lateral membranes

Anillin mutants additionally present with membrane degradation and nuclear displacement (Field et al., 2005). Depletion of anillin in *scra* RNAi gastrulas also results in the gradual disintegration of lateral membranes, which eventually form vesicles (Figure S5). Membrane degradation increases in severity over time. Some VF cells become multinucleated, and VF nuclei are intermittently displaced into the yolk sac (Figures 1H’,I’, Figure S5). These findings suggest that Anillin plays a role in VF formation – not only during actin ring closure and basal membrane formation, but also in maintaining membrane stability.

The possibility remained that the phenotypes seen in immunostainings reflected differences in the localization of our Neurotactin membrane marker, rather than degradation or loss of actual membranes. To substantiate our findings, we examined the fine structure of epithelial membranes using transmission electron microscopy (TEM) (Figure 2). At early stages of VF formation, interstitial spaces between ventral cells (indicating the presence of membranes) are visible in control embryos at low magnification (Figure 2A) indicating the presence of intact membranes (traced in green). In early stage *scra* RNAi gastrulas, these interstitial spaces are less intact, and lateral membranes have started disintegrating into vesicles (traced in red) (Figure 2B). Ventral cells in late stage control gastrulas have intact lateral and basal membranes (Figure 2C), all of which are absent in late stage *scra* RNAi gastrulas (Figure 2D). Specifically, at this late stage, interstitial spaces are no longer visible and have been replaced by an increased number of vesicles (Figure 2D); in other words, lateral membranes have been lost. Together, these data substantiate our findings with Neurotactin immunostaining, and indicate that basal membranes never form in *scra* RNAi embryos, and lateral membranes degrade late in VF formation.

**Figure 2.**
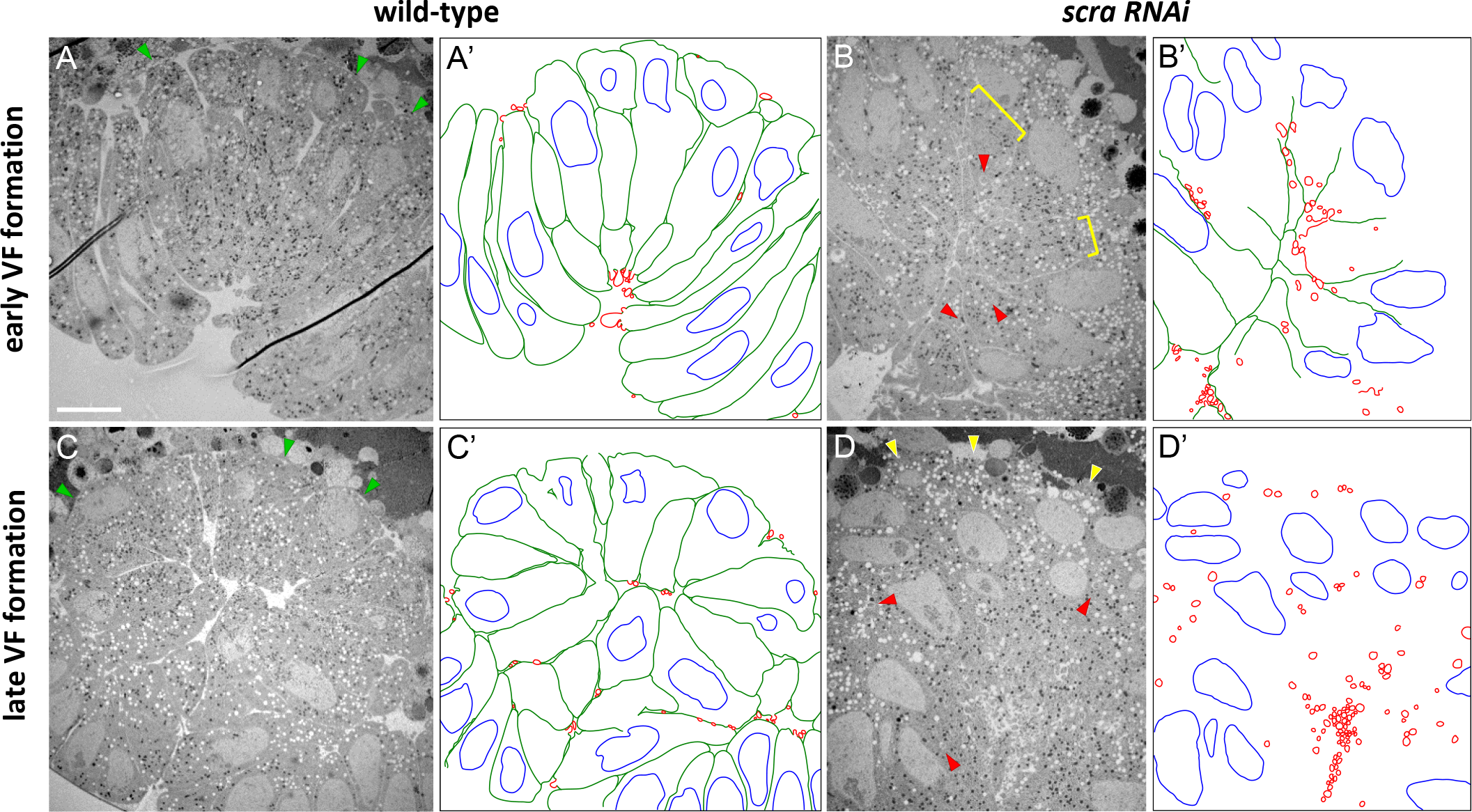
*scra* RNAi VFs remain folded despite degradation of lateral membranes (A-D) Low mag TEM of embryo sections prepared using a combined high pressure freezing/freeze substitution method during invagination (A,B) or late VF formation (C,D) in wild-type (A,C) and *scra* RNAi (B,D) embryos. Interstitial spaces are visible on the basal sides of VF cells (green arrows) in wild-type embryos during both early (A) and late (B) VF formation. Basal interstitial spaces are absent (yellow brackets) and lateral membranes are starting to degrade into vesicles (red arrows) in early VF formation *scra* RNAi embryos (B). Basal interstitial spaces are still absent (yellow arrows) during late VF formation in *scra* RNAi embryos (D) and lateral interstitial spaces are no longer visible, replaced by an increased number of vesicles (red arrows). (A’-D’) are hand-drawn traces of (A-D), respectively, highlighting intact interstitial spaces (green), nuclei (blue), and vesicles (red). Scale bar (10μm) applies to all images.

In spite of the gradual loss of membrane integrity during late stages of VF formation, the mesoderm nuclei maintain their ring-like spatial distribution. This indicates that the furrow retains its shape in the absence of basal, and subsequently lateral, membranes.

### VF formation without basal membranes can be explained by viscosity

Many existing models of VF formation require volume conservation within cells (Rauzi et al., 2013). It is unclear how such models could describe a mutant in which basal membranes are missing. A mechanistic question remains: how are *scra* RNAi embryos capable of forming a furrow while cells remain open?

To explore this question, we first developed a 2D model representing a transverse cross-section of the embryonic epithelium. In the model, the vitelline membrane is represented as a circular no-slip boundary encasing the model tissue. The perivitelline fluid, which fills the narrow space between the vitelline membrane and embryo, is modeled as an inviscid Newtonian fluid of fixed (low) viscosity. Cell membranes are modeled as series of short elastic springs. Note that when we refer to “membranes” in this section, we actually are referring to the membrane along with the associated load-bearing cytoskeleton. The yolk and cytoplasm within the embryo is modeled as a Newtonian fluid of viscosity η.

To model ventral furrow formation, we applied contractile stresses in a pattern chosen to mimic the forces thought to be present *in vivo* (Dawes-Hoang et al., 2005, Gracia et al., 2019, Leptin et al., 1992). Specifically, we assume that all membranes are subjected to a relatively small amount of constitutive stress, while the 16 mesodermal cells have additional stress applied to their apical and lateral membranes; this additional stress is linearly ramped up over time. Technical details of model implementation are described in the Supplemental Methods; the code used to generate the results is publicly available under https://github.com/doubrovinskilab/anillin_code.

In the version of the model representing wild type embryos, the mesoderm constricts apically and then invaginates as expected (Figure 3A). When basal membranes are removed, mesoderm invagination still occurs (Figure 3B), mimicking our observations in *scra* RNAi embryos. We had wondered how cytoplasmic contents remain within the interior of open cells in *scra* RNAi mutants; this model gave us an opportunity to explore the physical mechanisms underlying this phenomenon.

**Figure 3.**
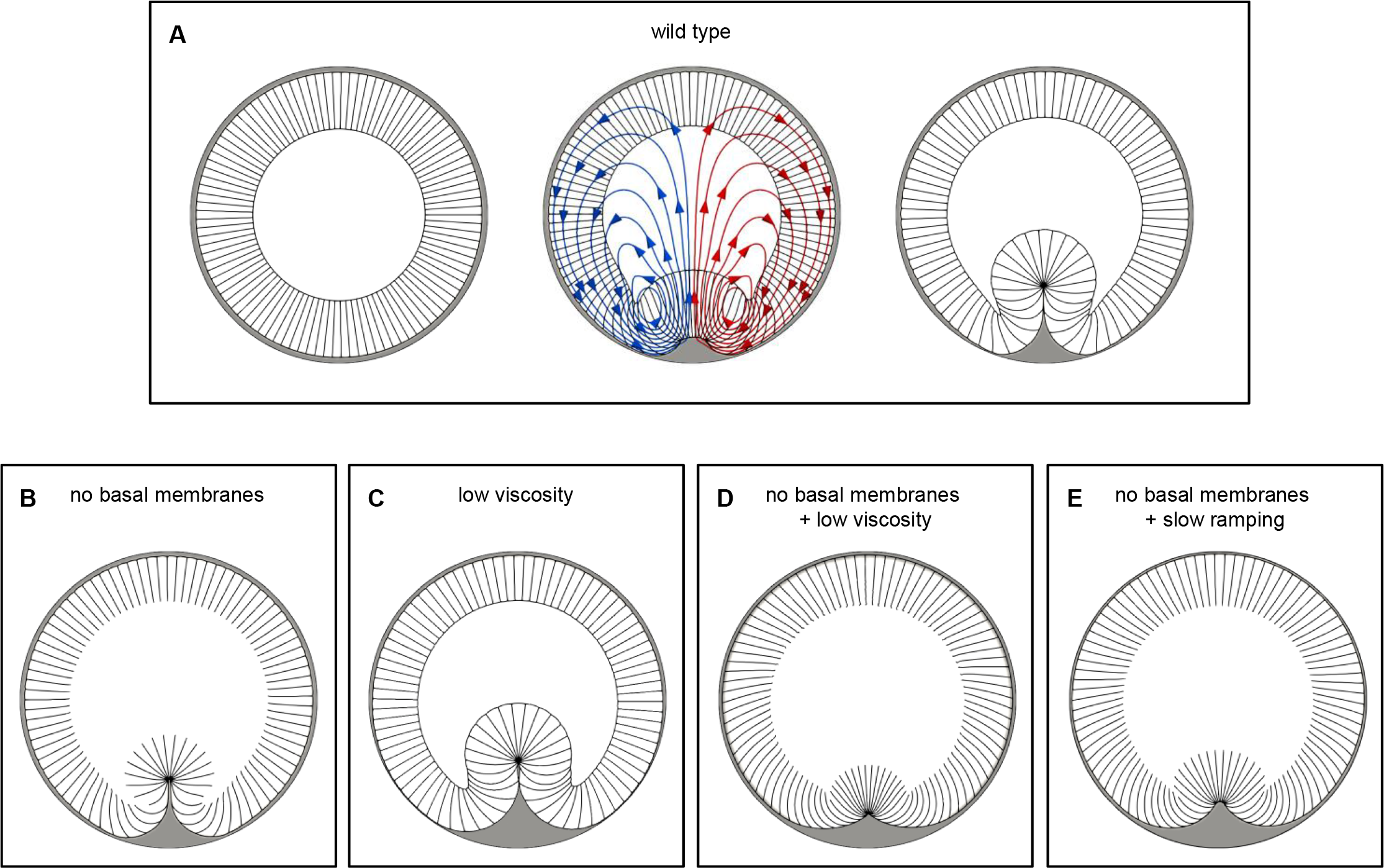
Simplified two-dimensional model of VF formation. (A) Two-dimensional model of VF formation. The initial state of the tissue is shown on the left. Middle panel shows a transient intermediate state of tissue during invagination; instantaneous fluid flow lines are also shown in red and blue. Right panel is the final state of the furrow (once maximal invagination depth has been reached, see Methods). (B)-(E) Final states of similar simulations in which specific material parameters or features have been altered. (B) Same parameters as in (A) without the basal membranes. Cytoplasm can move between the cellular interiors and the yolk sack unobstructed. VF still forms successfully. (C) Same parameters as in (A), except with the value of cytoplasmic/yolk viscosity reduced 100- fold. VF forms successfully. (D) Same parameters as in (A), except without the basal membranes and with the value of the viscosity reduced 100-fold. The depth of VF invagination is markedly reduced. (E) Same parameters as in (B) except that active stresses are ramped up 33-fold more slowly. The depth of VF invagination is markedly reduced. We conclude that VF invagination requires cell volume conservation either due to the presence of the basal membranes (as in the wildtype case), or due to the presence of sufficient viscous shear forces confining the cytoplasm to the cellular interior in the absence of the basal membranes (as in the *scra* RNAi background).

The reason we explicitly included fluid in our model is that we intuitively felt that shear (or viscous) forces may be responsible for invagination in *scra* RNAi mutants. Specifically, we hypothesized that shear forces imposed an effective volume constraint that prevented fluid from quickly leaving cells as they deformed. To test this hypothesis in our model, we reduced the value of η (the yolk and cytoplasmic viscosity) 100-fold. With basal membranes present, apical constriction and mesoderm invagination occur normally (Figure 3C), but in the absence of basal membranes, mesoderm invagination is greatly reduced (Figure 3D). Furthermore, this reduction is accompanied by a shortening of lateral membranes and an apparent reduction of cytoplasmic volume within the mesoderm. These results support our hypothesis that shear forces play a key role in maintaining some degree of volume conservation in partially open cells during this morphogenetic movement.

Shear forces are a product of two factors: the viscosity of the fluid, and the velocity gradient (which in the case of fluid moving between two surfaces is proportional to velocity itself). We therefore next examined the effects of changing the rate at which we ramped up contractile stresses in the mesoderm, since slower ramping would be expected to yield lower velocities and thus lower shear forces. We decreased the ramping speed 33-fold and allowed the simulation to run longer (to reach the same final values for stress); this again resulted in greatly reduced mesoderm invagination (Figure 3E). This result confirmed that shear forces are required for invagination in models with basally open cells.

Importantly, these results also show that the quantitative details included this model affect the qualitative outcome: changing either the viscosity of the fluid or the ramping rate of the applied stresses can greatly affect the extent of invagination. This highlights the importance of considering a realistic, physically accurate model if we want use it to explore physical effects that are actually relevant to the biological system.

The 2D model considered above is useful for exploring general physical effects, but is more toy model than realistic representation. In particular, the parameters used were not based on measured values and were in fact in arbitrary units.

However, we have recently developed a physically realistic 3D model of the *Drosophila* embryo which incorporates experimentally measured geometries and material properties (Cheikh et al., 2022). In particular, the cytoplasmic and yolk viscosity, and the Young’s (elastic) modulus of the apical and basal-lateral membranes were all measured or estimated from experimental data (Cheikh et al., 2022, Doubrovinski et al., 2017) (see also methods section). To address whether shear stresses are relevant to ventral furrow formation *in vivo*, we therefore asked to what extent shear stresses affect ventral furrow formation using this biophysically realistic model.

To study VF formation in this 3D model, we again linearly ramped up apical and lateral membrane stress over time in the mesodermal cells. For a range of ramping speeds, mesoderm invagination appeared relatively normal in cases both with basal membranes (equivalent to wild type) and without basal membranes (equivalent to *scra* RNAi mutants) (Figure 4). However, ramping speeds did affect the depth of furrow formation in the *scra* RNAi mutant case (Figure 4D). For example, a ramping speed that reached maximal furrow depth around 1 minute (Figure 4B) had a furrow depth of 33 microns, while a 3-fold slower ramping speed resulted in a final depth of 26 microns at around 3 minutes (Figure 4C).

**Figure 4.**
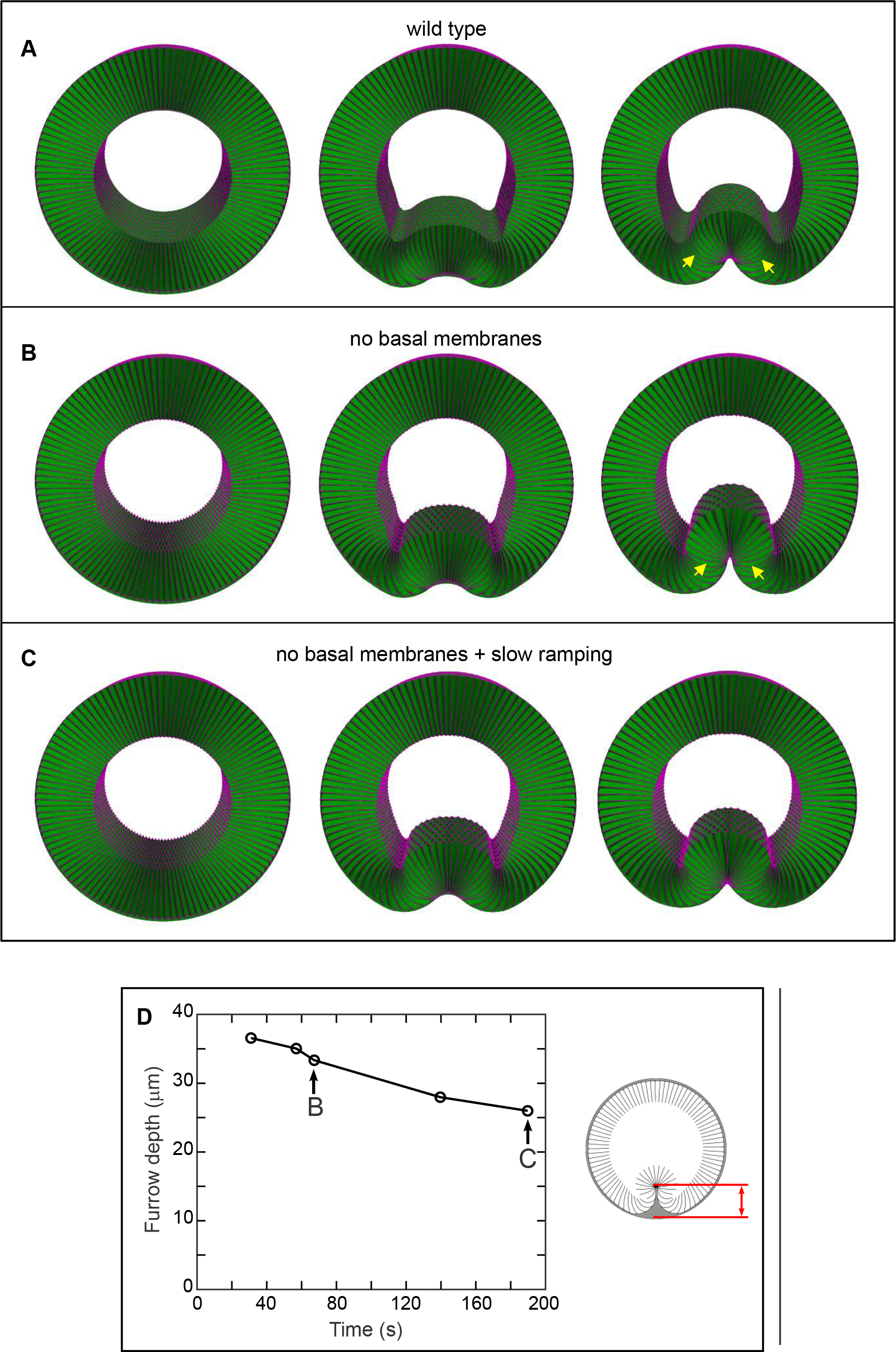
Three-dimensional model of VF formation based on *in vivo* measurements. (A) VF formation with basal membranes present (wildtype case). Left panel is starting configuration, middle panel is an intermediate time point, and right panel is the final state of the furrow (once maximal invagination depth has been reached, see Methods). (B) Same as (A) without the basal membranes. The VF forms successfully. Cells flanking the mesoderm (arrows) are significantly shorter than the corresponding cells in (A) (quantified in Supplemental Figure S6). (C) Same as (B) except that active stresses are ramped up 3-fold more slowly. Maximal invagination is reached approximately 3 min into force ramping (compared to approx. 1 min in both (A) and (B)). (D) Simulations of tissue without basal membranes were run using a variety of ramping speeds. For each simulation, the maximal furrow depth (see red arrow in schematic on the right) is plotted as a function of the time taken to reach maximal VF invagination. Indicated points correspond to the simulations shown in (B) and (C).

*In vivo*, ventral furrow formation occurs over approximately 5 to 10 minutes (Gelbart et al., 2012). This is slightly longer than the time scale on which we start to see decreasing maximal furrow depths in our model. This could indicate that the parameter values we used may be slightly off, for example up to about 2-fold; this is in fact consistent with the expected error range of our measurements (Cheikh et al., 2022). Alternatively, this could indicate that there are additional contributing factors that help allow ventral furrow formation to proceed normally in *scra* RNAi mutants. One potential such factor might be cell rearrangements, which we do not include in our model. Allowing cell rearrangements would effectively decrease the elasticity of the apical surface, which could allow for deeper furrow invagination. Although we believe such corrections could yield an even more realistic model, we hold that the close match between experimental results and our 3D model strongly suggest that shear stresses play a major role in ventral furrow morphogenesis.

## DISCUSSION

In this work, we explored the role of basal membranes in epithelial invagination by examining VF formation in *scra* RNAi embryos, in which basal membranes do not form. Invagination proceeded normally, which was surprising in light of previous computational models which posited that cellular volume conservation is required for VF invagination (Gelbart et al., 2012, Rauzi et al., 2013). Using both a 2D toy model and a biophysically realistic 3D model with experimentally-derived parameters, we explored the physical mechanisms underlying this surprising result. This analysis provides strong evidence that viscous shear is a key mechanical effect contributing to VF morphogenesis. In other words, VF formation may be considered a “swimming phenomenon” where solid structures (cellular membranes) move by exerting force against the ambient fluid.

Several additional conclusions can be drawn based on our study.

Our biophysical model shows that the time scale of tissue invagination is determined by the rate with which active stresses (for example, from myosin accumulation) build up, and not by the time scale of the resulting viscoelastic response. This is equivalent to stating that this tissue deformation occurs adiabatically, meaning that the system is in mechanical equilibrium at all time points. This conclusion is also supported by previously described direct physical measurements (Cheikh et al., 2022).

Previously published analyses of VF formation indicate that mesodermal cell elongation precedes invagination (Sweeton et al., 1991), so these are distinct processes. Our analysis here shows that cells elongate even in *scra* RNAi embryos that don’t have basal membranes at all (Figure 1H). Therefore, cell elongation, like the later process of invagination, does not require pressure difference across the basal membranes, and can be driven by shear forces.

Importantly, our computational model indicates that basal membranes and viscous shear forces play complementary mechanical roles in wild type embryos, since either is sufficient for these morphogenetic behaviors. In this particular system, the importance of viscous shear forces was only revealed experimentally in the *scra* RNAi background. However, there are many tissues in which similar effects might play the primary role. In the housefly *Musca vicina*, for example, it has been reported that gastrulation proceeds while the blastoderm in still syncytial (Bhuiyan and Shafiq, 1959). Additionally, phenomena involving viscous flows have been unexpectedly implicated in *Drosophila* oocyte growth (Lu et al., 2022). Further afield evolutionarily, we note that for example the slime mold *Physarum polycephalum* (Alim et al., 2017) and the glass sponges (class *Hexactinellida*) (Leys et al., 2006) generate complex 3D shapes as syncytial organisms, suggesting that shear forces should be investigated in these model systems as well.

## METHODS

### Drosophila genetics

For anillin depletion experiments (Figures 1-2), we used the following genotype: UAS-Nrt-eGFP; P{y[+t7.7] v[+t1.8]=TRiP.GL01269}attP2/mat15-GAL4. TRiP.GL01269 was derived from RRID:BDSC 41841 (Bloomington Drosophila Stock Center); mat15-GAL4 was derived from RRID:BDS_80361 (Bloomington Drosophila Stock Center). To generate UAS-Nrt-eGFP, full-length Nrt-RB was cloned by PCR from LD22004 (Drosophila Genomics Resource Center stock #5736) and inserted into pPWG (Drosophila Genomics Resource Center stock #1078).

This plasmid was injected by BestGene using P-element insertion.

### Fluorescent immunohistochemistry

Embryos were collected from female progeny on grape agar plates supplemented with yeast paste after ≥3.5hrs. Fly embryos were heat-methanol fixed as described previously (Muller and Wieschaus, 1996). The block, primary antibody, and secondary antibody steps were all done overnight with nutation at 4°C. Primary and secondary antibodies were diluted in block solution (1x PBS, 0.1% Triton X-100, and 5% heat inactivated goat serum or 0.2% BSA). Antibodies used include mouse anti-Neurotactin (1:50) (DSHB), guinea pig anti-Snail (1:2000) (gift from Eric Wieschaus), goat anti-mouse IgG-Alexa Fluor 488 (1:500) (Invitrogen), and goat anti-guinea pig IgG-Alexa Fluor 568 (1:500) (Invitrogen). Nuclear staining was done using DAPI (1μg/mL) (Invitrogen).

### Confocal fluorescence imaging of embryo sections

Immunostained fly embryos were staged in 1x PBS with 0.1% Triton X-100 under bright field on an Accu-Scope dissection microscope. A coverslip was prepared by cutting off ∼1cm of the short edge and placing it in the center of the coverslip, perpendicular to the long edge, then adding a linear pool of AquaPolymount along the supporting glass strip. Selected gastrulas were transferred to the pool of mounting medium and sectioned along the dorsal-ventral axis using a 22g needle. Embryo halves were positioned cut side down, leaning against the supporting glass strip. Z-stacks were imaged using a Plan-Apochromat 63x/1.40 Oil objective on a Zeiss LSM 700 confocal microscope. For immunohistochemistry experiments, we fixed and stained samples over the course of several weeks, until we had at least 40 separate gastrulas imaged. From these, we selected all samples in which the morphology had not been significantly distorted by sectioning damage or angle – these results are quantified in the Figure 1 bar graphs.

### Statistical analysis of immunostaining results

For the purposes of all experiments, all samples are biological replicates. We binned data from bar charts in Figure 1 (19 positions, -9 to 9) in sequential groups of 3, producing a plot with 6 average values of membrane lengths along the VF. We then compared experimental groups pairwise using Fisher’s linear discriminant analysis. Specifically, we produced two sets of projections on the direction normal to the discriminant hyperplane, each set corresponding to one of the experimental groups being compared. Finally, the significance of the difference between the two sets of projections was assayed using the pairwise Kolmogorov-Smirnov test. An exception was made in the case of late VF formation *scra* RNAi embryos, where we only binned data from 15 positions (-7 to 7) – this was due to a smaller sample size for this group. For Figure S4, all data were used in analysis.

### Transmission electron microscopy

Briefly, we froze gastrulas of various stages under high pressure, subjected them to freeze substitution, made 60-70nm transverse cross sections, and imaged them under TEM. A combined high pressure freezing and freeze substitution (HPF/FS) method was used to fix fly embryos as described elsewhere (Zhang and Chen, 2008). A Wohlwend Compact 03 high pressure freezer was used to fix the embryos. Samples were freeze substituted in 1% osmium tetroxide, 0.1% uranyl acetate in 98% acetone and 1% methanol using a Leica EM AFS2. The embryos were embedded in Embed-812 resin and polymerized in a 60°C oven overnight. Blocks were sectioned with a diamond knife (Diatome) on a Leica Ultracut 7 ultramicrotome (Leica Microsystems) and collected onto slot grids and post stained with 2% aqueous uranyl acetate and lead citrate. Images were acquired on a JEOL JEM-1400 Plus TEM equipped with a LaB_6_ source operated at 120 kV using an AMT-BioSprint 16M CCD camera. For TEM experiments, we continuously staged and fixed embryos over the course of 6-9 hours. Embryo sections shown in Figure 2 are representative of 5 (Figure 2A), 3 (Figure 2B), 11 (Figure 2C), or 4 (Figure 2D) samples. From these, we selected all samples in which the morphology had not been significantly distorted by ice/mechanical damage or other staining artifacts.

### Mathematical modeling

For a detailed description of computational methods used, please see the Appendix.

## ACKNOWLEDGEMENTS

This work was supported by the Welch Foundation award I-1950-20180324 and the National Institutes of Health grant 1R01GM134207-01. Stocks obtained from the Bloomington Drosophila Stock Center (NIH P40OD018537) were used in this study. The authors would like to acknowledge the assistance of the UT Southwestern Electron Microscopy Core, especially Anza Darehshouri. The NIH Shared Instrumentation award 1S10OD021685-01A1 to Katherine Luby-Phelps allowed for use of the JEOL JEM-1400 Plus TEM.

## COMPETING INTERESTS

The authors declare no competing interests.

## SUPPLEMENTAL FIGURES

**Figure S1.**
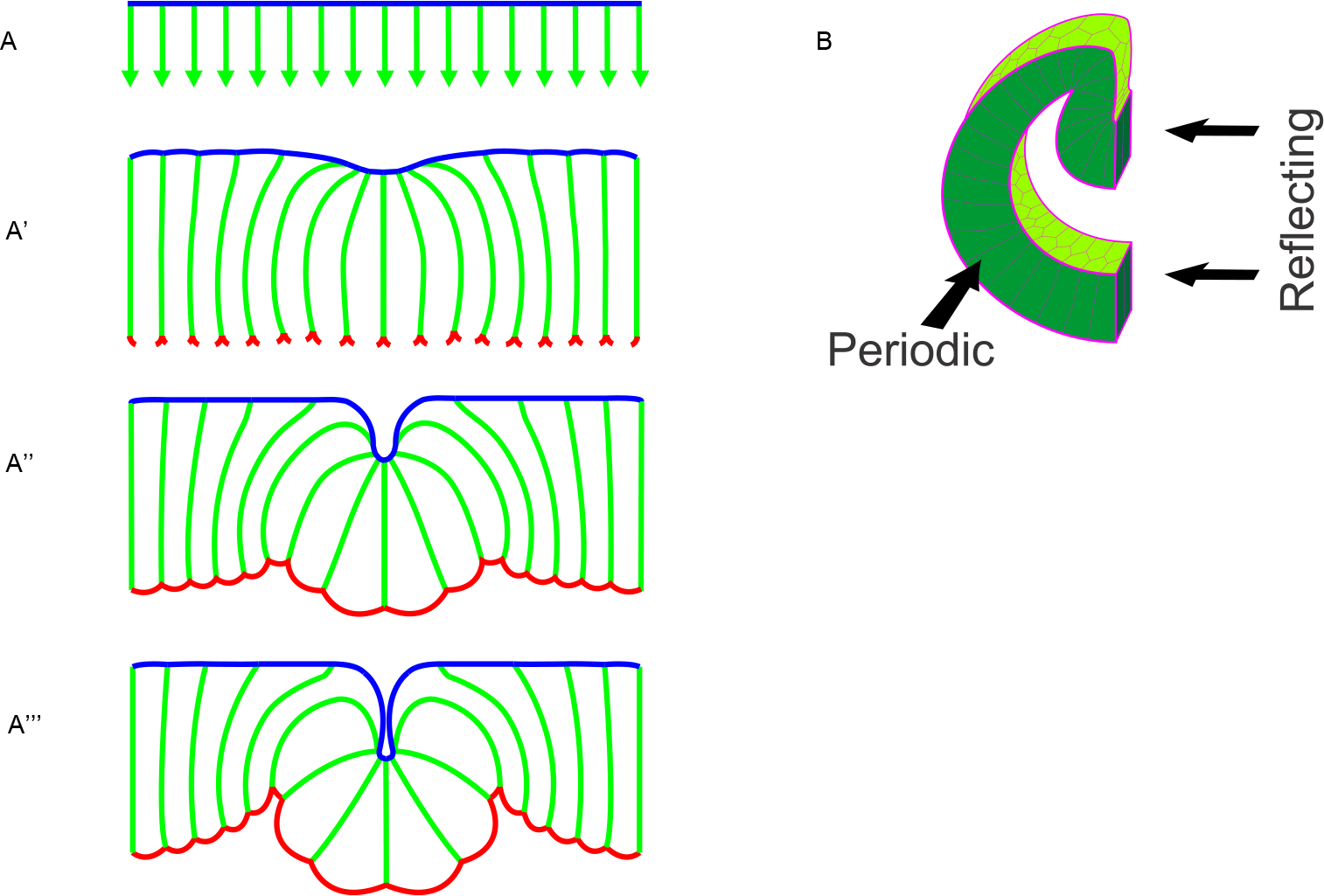
Schematic of membrane shape changes throughout VF formation. (A-D) Illustration of VF formation in cross section showing shape changes in apical (blue), lateral (green), and basal (red) membranes throughout tissue folding. Lateral membranes lengthen (arrows) throughout cellularization (A). Lateral membranes of VF cells lengthen more as apical membranes constrict to initiate VF formation (B). Lateral membranes shrink back down as tissue invaginates and basal membranes form. Basal membranes seal and expand

**Figure S2.**
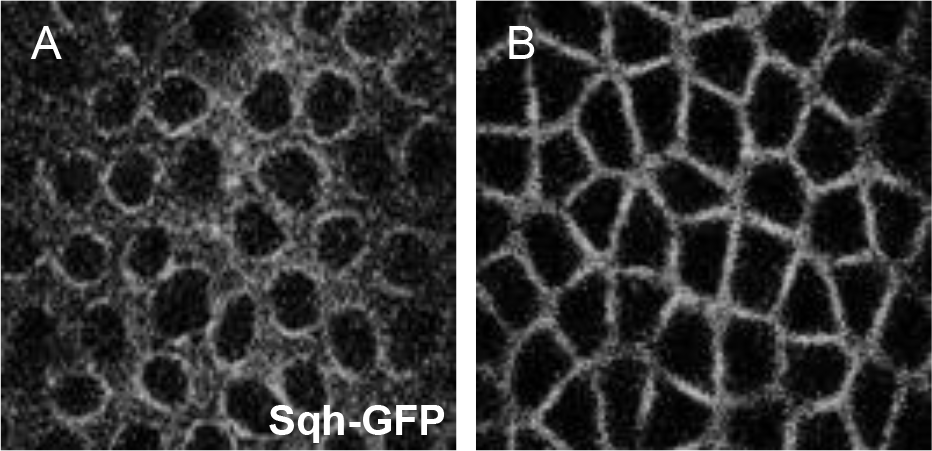
Acto-myosin ring closure is delayed in *scra* RNAi embryos. (A,B) Maximum intensity projections of the cellularizing ventral epithelium in live control (A) and *scra* RNAi (B) embryos during late cellularization. Acto-myosin rings remain open and are misshapen in *scra* RNAi embryos, preventing the formation of basal membranes.

**Figure S3.**
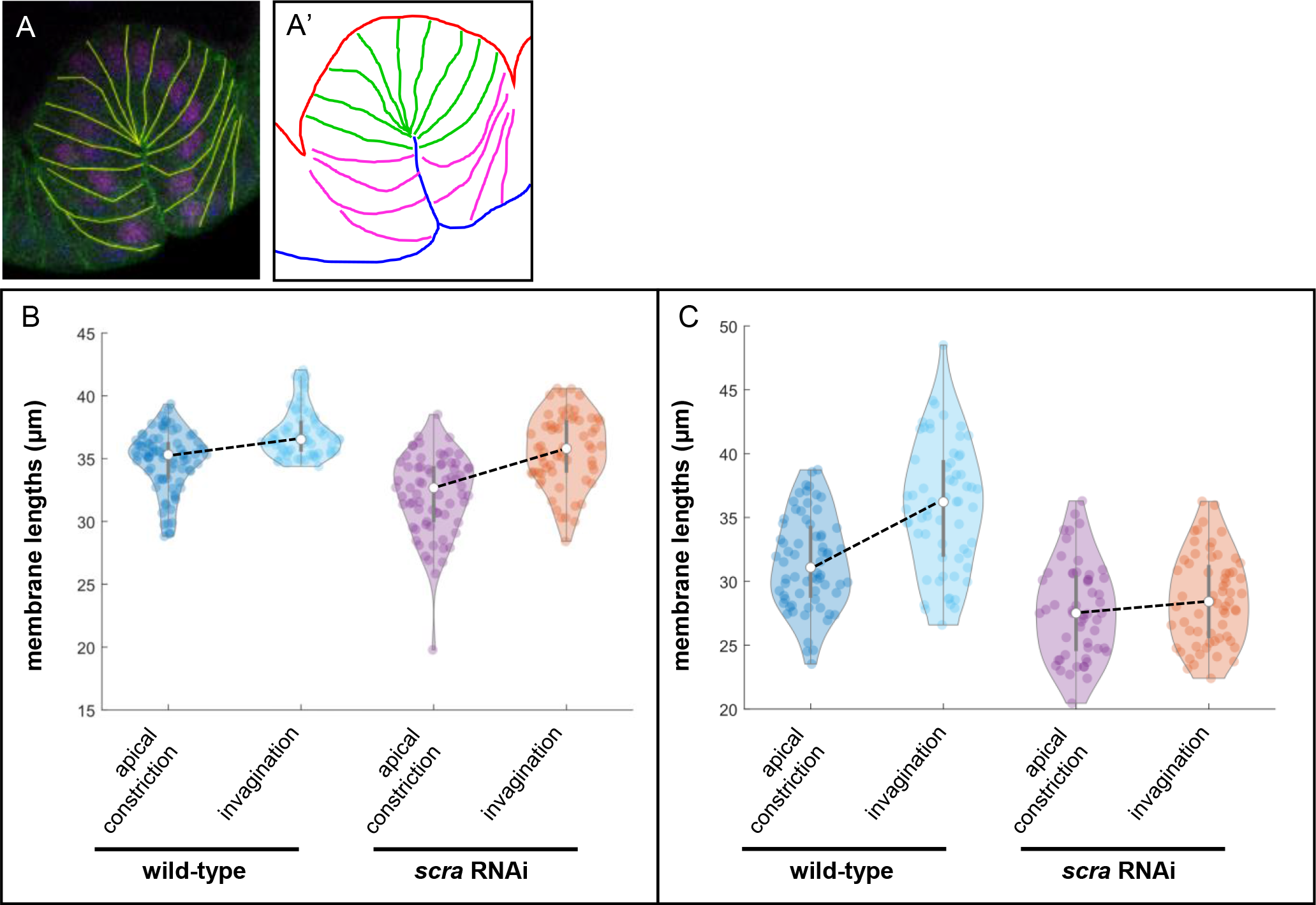
Central mesodermal cells lengthen more, peripheral mesodermal cells shorten less throughout VF formation in *scra* RNAi embryos compared to wild-type. Example of designated central (green) and peripheral (pink) mesoderm regions shown in (A,A’). Violin plots show changes in average central (B) and peripheral mesodermal membrane lengths (C) between apical constriction (ac) and invagination in wild-type (n_ac_ = 9 embryos, n_invag_ = 7 embryos) and *scra* RNAi embryos (n_ac_ = n_invag_ = 8 embryos).

**Figure S4.**
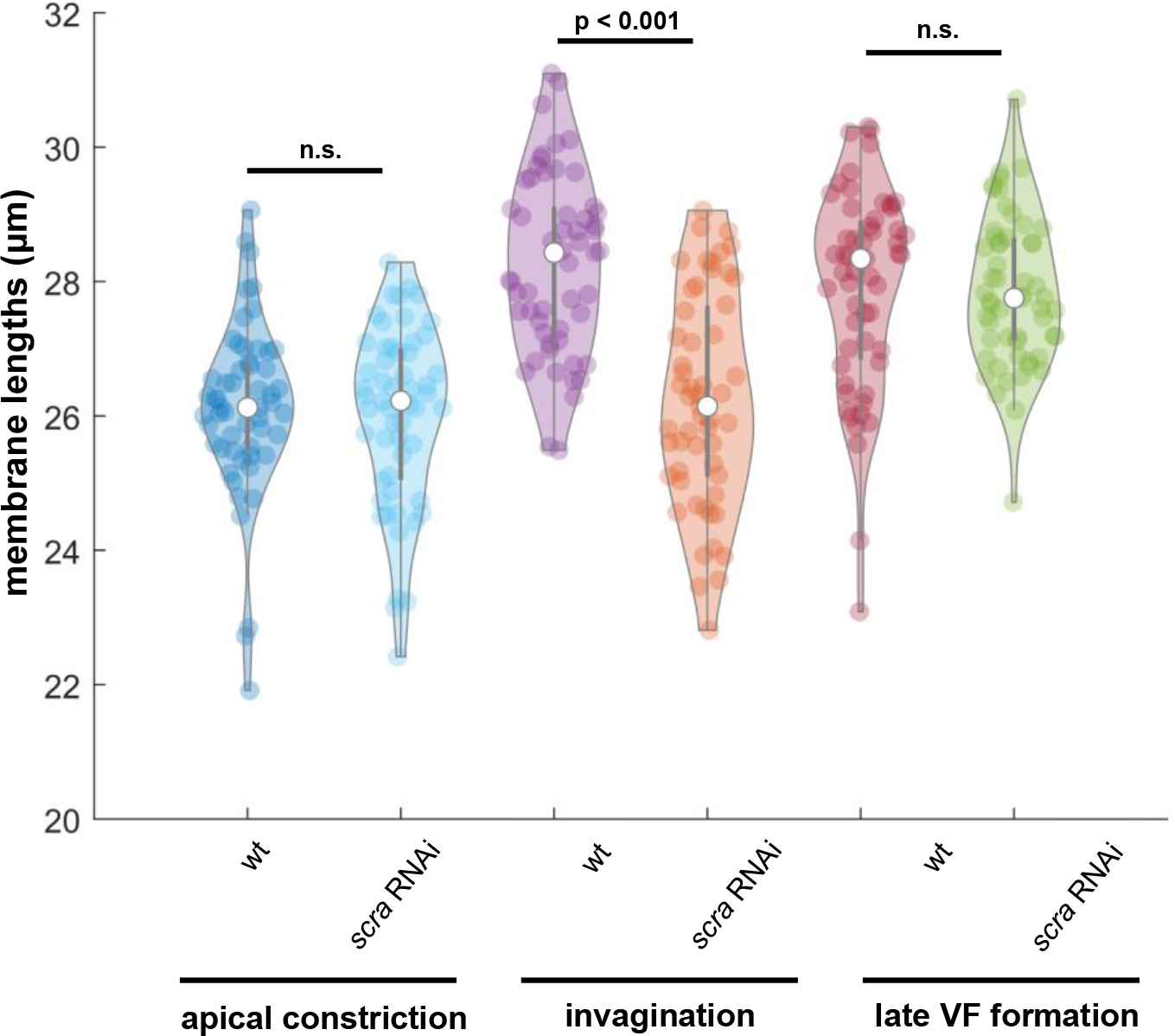
Non-mesodermal cells are shorter during invagination in *scra* RNAi embryos. Violin plots of non-mesodermal cell lengths during apical constriction, invagination, and late VF formation in wild-type (wt) and *scra* RNAi embryos. Non-mesodermal cells of *scra* RNAi embryos do not lengthen as quickly as those in control, resulting in shorter epithelium mid VF-formation. By late VF formation, non-mesodermal cells of *scra* RNAi embryos have reached lengths similar to those in wild-type embryos.

**Figure S5.**
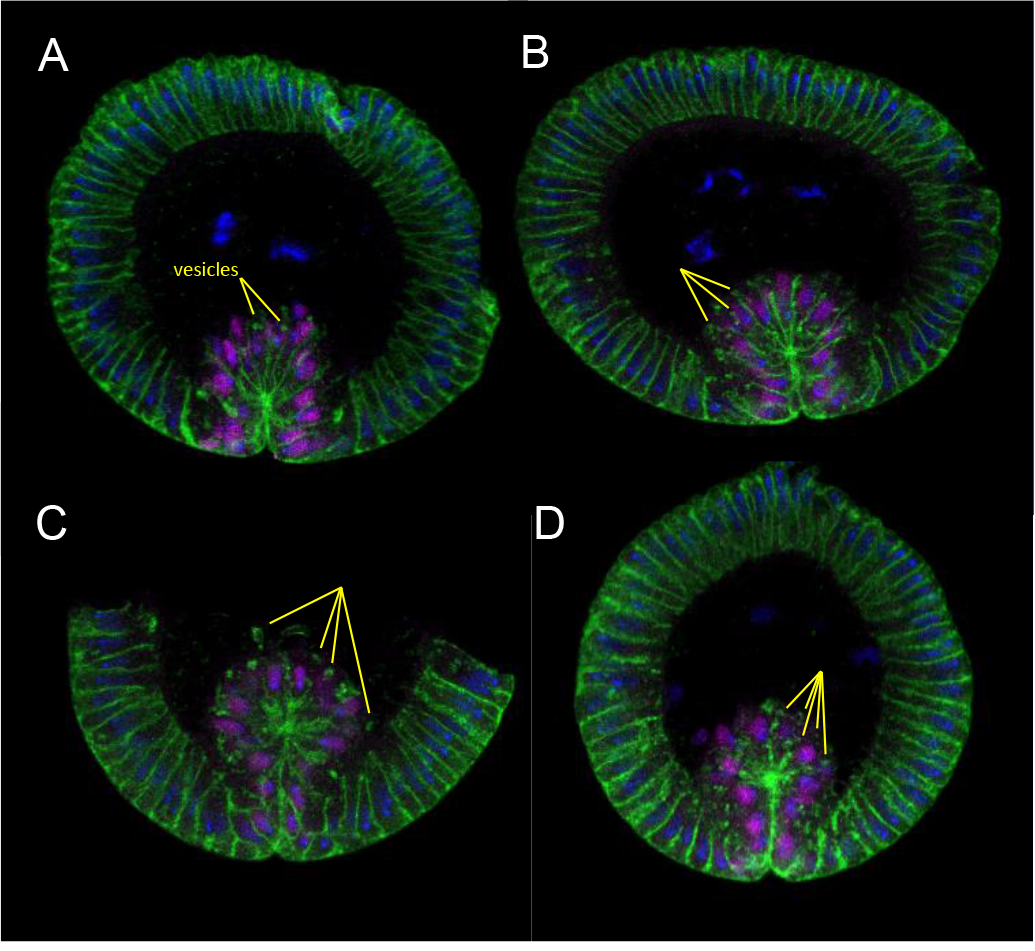
Membranes in *scra* RNAi embryos deteriorate more over time. Confocal immunofluorescence of heat/methanol-fixed anillin RNAi embryos at increasingly older stages of gastrulation (A-D). Membrane degradation increases with age, and vesicles become more numerous. Note: the embryo section in C is incomplete due to how it was cut, not any depleted protein expression or lack of antibody staining.

**Figure S6.**
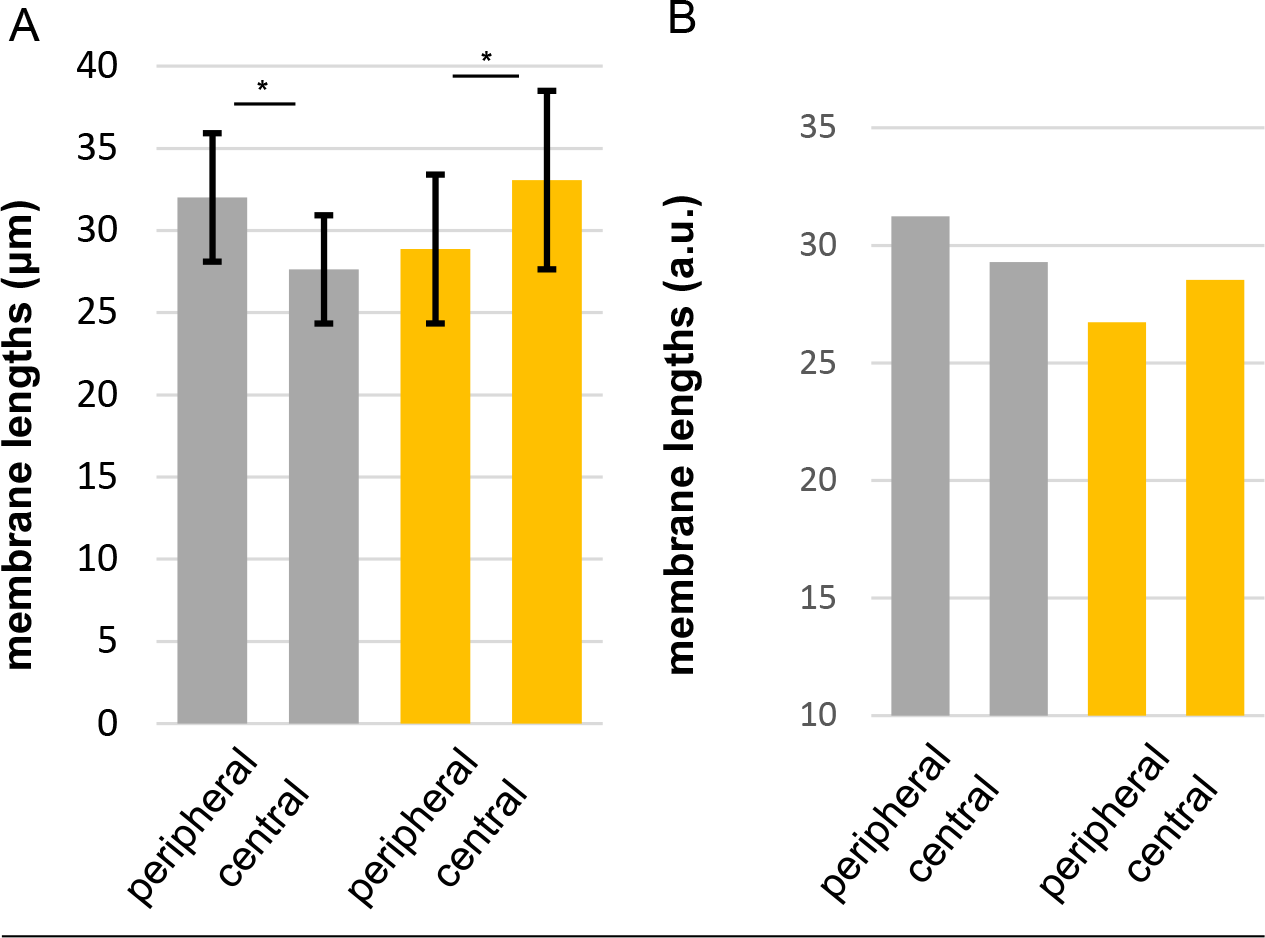
Model accounts for *in vivo* membrane length differences between membranes at edge and center of mesoderm. (A) Lengths of membranes in peripheral VF compared to central VF membrane (see figure S3A) in control (grey, n = 14 embryos) and *scra* RNAi (yellow, n = 7 embryos) embryos during late VF formation. Data are represented as mean ± s.d. Significance was determined by two-way Kolmogorov-Smirnov test (* = p < 0.05). (B) Lengths of equivalently- located membranes in 3D model control (grey, Figure 3I) and 3D model *scra* RNAi (yellow, Figure 3L) embryos at an equivalent stage.

## APPENDIX

### Numerical simulations of ventral furrow formation

In this section, we detail the numerical method used to simulate ventral furrow formation. Our approach is a combination of the Immersed Boundary Method (IBM) (Peskin, 1972; Peskin and Printz, 1993) and the Finite Element Method (FEM) (Boffi et al., 2007). The choice of the numerical method is motivated by the requirement to describe fluid-structure interactions, since epithelial cells comprising gastrulation have previously been shown to be predominantly elastic on their surface and viscous in the interior (Cheikh et al., 2022; Doubrovinski et al., 2017; Rauzi et al., 2008). A key advantage of the IBM approach is that the computational (Eulerian) grid representing the fluid remains fixed throughout a simulation, while the solid (Lagrangian) mesh deforms. This approach avoids the need to re-mesh the fluid throughout the simulations. Technical details of our numerical implementation and validation of the method are given elsewhere (Cheikh et al., 2022).

### Governing Equations

For both 2D and 3D simulations, time evolution of the fluid-immersed interfaces representing cellular edges requires simulating the ambient fluid, whose dynamics are given by the Stokes equations:

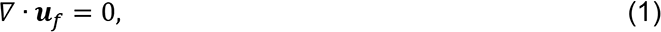

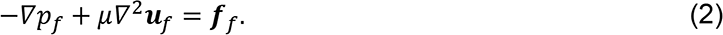

Here *μ* is dynamic viscosity, while *u*_*f*_(*x,t*) and *p*_*f*_(*x,t*) are the velocity and the pressure fields respectively. *f*_*f*_ is the force density acting on the fluid. Importantly, *f*_*f*_ includes the force on the fluid from the immersed structure. We are justified in neglecting inertia since a typical Reynolds number in our problem may be estimated to be on the order of Re = 10^−8^ (Doubrovinski et al., 2017; Selvaggi et al., 2018). Cell edges are described as linearly elastic shells characterized by a (two-dimensional) Young’s modulus *E* and Poisson’s ratio σ. As detailed in (Seung and Nelson, 1988), it is convenient to approximate the immersed solid surfaces as a triangulated mesh of approximately equilateral triangular elements. Setting the spring constant of triangle element edges to *k*, in the continuum limit one recovers an approximation of a linearly elastic sheet with a Young’s modulus of 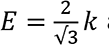 and a Poisson’s ration of 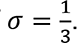 The dynamics are driven by active compressive stresses (corresponding to the action of myosin motors) described as force dipoles of constant magnitude (i.e. independent of the length of an edge), directed along the edges of the triangular elements discretizing the solid. In this way, the *i*^*th*^ solid edge contributes a force dipole to the *f*_*f*_-term on the right-hand side of equation (2):

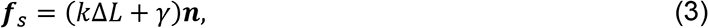

where *k* is the elastic constant of a solid edge, Δ*L* is elongation distance, *γ* determines the magnitude of active stress, and *n* is the unit vector along the edge.

The dynamics of the solid are coupled to that of the fluid by transferring force density from solid nodes to the adjacent fluid nodes and advecting the solid with the local velocity of the fluid by using the “spreading” and the “interpolation” of the standard IBM (Boffi et al., 2007). Validation of the numerical scheme and further technical details of our implementation are given in (Cheikh et al., 2022). All simulation code is publicly available at the Doubrovinski lab GitHub link https://github.com/doubrovinskilab/anillin_code.

### The geometry of the 3D model

The geometry of the model is illustrated in Figure 4. To reduce computational time, we take advantage of reflection symmetry of the embryo and only simulate left half of gastrula (by imposing reflecting boundary condition along the dorso-ventral axis as well, see Figure 4). We also take advantage of the fact that the cross-sectional shape of the embryo during ventral furrow formation is constant for most of the length of the embryo, varying only at the round poles. Therefore, we simulate what is essentially a cylindrical embryo by simulating a one-cell-thick cross-section with periodic boundary conditions on both cross-sectional faces. The outer solid boundary represents the rigid vitelline membrane where no-slip boundary condition is imposed. The cellular layer is separated from the model vitelline membrane by a thin layer of perivitelline fluid whose viscosity is set to that of water. Cells are hexagonal prisms whose faces are elastic sheets modelled as a set of inter- connected springs as described in the previous section. Cytoplasm and yolk are both modelled as Newtonian fluid with a viscosity of 1000 cP, as previously measured (Doubrovinski et al., 2017; Selvaggi et al., 2018). The dimensions of cells and their geometries are taken directly from experimental measurements.

Elasticities of cellular edges are taken from measurement data in our recent work where cellular edges of a developing embryo were directly probed mechanically, using bendable cantilevers (Cheikh et al., 2022). In fact, the model used in the present paper is the same as that described in (Cheikh et al., 2022), except for two additional features. (1) Active forces are imposed on cellular edges to drive tissue folding and (2) basal elasticity in this model is set equal to the lateral rather than basal elasticity from our model in (Cheikh et al., 2022). The reason for this choice is that the previous model described a cellularizing embryo where the basal surface consists of a hexagonal grid of cell edges that are highly enriched with actin. During cellularization, these hexagons shrink until they close off the basal sides of the cells, at which point the enriched actin disappears. Therefore we expect the basal surfaces during ventral furrow formation to be molecularly and mechanically equivalent to the lateral rather than basal surfaces during cellularization.

### Simplified (2D) model

The computational approach used for the simplified two-dimensional simulations was completely analogous to the one used in 3D simulations, but the geometrical and parameters were chosen differently for simplicity. The computational domain was chosen as a disc representing a cross section through the embryo, with no-slip condition imposed at the boundary (the model vitelline membrane. Membrane elasticity was assumed to be the same in all cellular domains and uniform throughout the embryo. The interior of the embryo (cytoplasm and yolk compartments) was described as a Newtonian fluid of fixed viscosity. Initially, the surface of the embryo is separated from the model vitelline membrane by a gap of uniform thickness (perivitelline space) filled with model perivitelline fluid, described as a relatively inviscid Newtonian fluid. Contractile stresses were imposed on a subset of membranes within the mesoderm (both apical and lateral). Simulations were done using a combination of Finite Element Method and Immersed Boundary Method, completely analogously to the 3D case, except adopting the methods to the two-dimensional geometry in the obvious way. Model parameters used in the different simulations shown in Figure 3 follow; we list these parameters in arbitrary units since they don’t correspond to the actual measured values. Simulation of the “wild type” embryo in Figure 3a: radius of the embryo was 117, thickness of the peri-vitelline space is 3. The embryo comprised 101 initially identical cells of height 35. The viscosity of the cytoplasm was 1, viscosity of the perivitelline space was 0.01. The elasticity of springs discretizing cellular boundaries was 0.008. “Mesoderm” was assumed to comprise 17 cells whose “apical” and “lateral” membranes were subjected to active stresses (constant force dipoles as was described for the 3D case). The ratio of apical to lateral active stresses in the mesoderm was fixed to 12.

Active stresses were ramped linearly according to 3.3·10^-8^ t + 2·10^-5^ (in the apical sides of mesodermal cells; stresses in the lateral membranes of the mesoderm had a 12-fold smaller value). All parameters in simulations from Figure 3B-E are the same as in Figure 3A except for the changes listed in the corresponding figure caption. All simulation code is available under https://github.com/doubrovinskilab/anillin_code.

## Notes

### Competing Interest Statement

The authors have declared no competing interest.

### Summary of Updates

The article includes quantitative 3D modeling entirely based on the work in https://www.biorxiv.org/content/10.1101/2022.08.12.503803v2.full This additional work establishes model parameters that are then used to test the proposed mechanism of tissue invagination quantitatively.

